# Multimodal evidence for delayed fear extinction in adolescence and young adulthood

**DOI:** 10.1101/355503

**Authors:** Jayne Morriss, Anastasia Christakou, Carien M. van Reekum

**Affiliations:** Centre for Integrative Neuroscience and Neurodynamics, School of Psychology and Clinical Language Sciences, University of Reading Reading UK

**Keywords:** Fear Extinction, Anxiety, Adolescence, Amygdala, mPFC

## Abstract

Previous research in rodents and humans points to an evolutionarily conserved profile of blunted fear extinction during adolescence, underpinned by brain structures such as the amygdala and medial prefrontal cortex (mPFC). In this study, we examine age-related effects on the function and structural connectivity of this system in fear extinction in adolescence and young adulthood. Younger age was associated with greater amygdala activity and delayed mPFC engagement to learned threat cues as compared to safety cues. Furthermore, greater structural integrity of the uncinate fasciculus, a white matter tract that connects the amygdala and mPFC, mediated the relationship between age and acceleration of mPFC engagement during extinction. These findings suggest that age-related changes in the structure and function of amygdala-mPFC circuitry may underlie the protracted maturation of fear regulatory processes, rendering younger individuals more vulnerable to anxiety disorders, which emerge during development.

## 1.1 Introduction

The ability to discriminate between cues that signal fear and safety is crucial to an organism’s wellbeing. This ability not only supports escape and avoidance defensive mechanisms, but also prevents disproportionate reactivity, ensuring long term protection against chronic stress and psychopathology. Brain circuitry at the core of fear discrimination learning includes areas within the amygdala and medial prefrontal cortex (mPFC) (Milad & Quirk, 2012). During fear acquisition, heightened amygdala activity is observed in response to a previously neutral cue that, through conditioning, comes to be associated with an aversive outcome (thereby becoming a conditioned stimulus, or CS) (Büchel, Morris, Dolan, & Friston, 1998). Subsequent fear extinction training, which involves repeated presentations of the CS without the aversive outcome, results in reduced amygdala responses to the CS over time (LaBar, Gatenby, Gore, LeDoux, & Phelps, 1998), with the mPFC playing a critical role (Milad & Quirk, 2012; Milad et al., 2007; Phelps, Delgado, Nearing, & LeDoux, 2004).

Recent research with rodents and humans has shown distinctive developmental profiles of blunted fear extinction in adolescents (Kim, Hamlin, & Richardson, 2009; Kim, Li, & Richardson, 2011; McCallum, Kim, & Richardson, 2010; Pattwell, Bath, Casey, Ninan, & Lee, 2011; Pattwell et al., 2012), such that adolescents continue to show signs of defensive responding to previously learned threat cues (freezing in rodents and elevated skin conductance in humans), suggesting an age-specific relative dysregulation of the process that updates threat associations as safe (Kim, Li, & Richardson, 2011; Pattwell et al., 2012). Immunohistochemical evidence in rodents points to reduced synaptic plasticity in areas of mPFC during adolescence (Kim, Li, & Richardson, 2011; Pattwell et al., 2012). However, direct evidence regarding the neural circuitry underlying blunted fear extinction in adolescent humans is lacking.

Immature amygdala-prefrontal cortical interactions are thought to be responsible for this blunted fear extinction seen in adolescence (Nelson, Lau, & Jarcho, 2014). The amygdala and mPFC undergo substantial structural change during development: the amygdala shows linear increases in grey matter across late childhood and adolescence, while the mPFC is characterised by quadratic changes, with substantial grey matter growth across childhood followed by grey matter pruning across adolescence (Gogtay et al., 2004; Østby et al., 2009; Wierenga et al., 2014). Furthermore, the uncinate fasciculus, a white matter tract that connects the amygdala and mPFC, undergoes protracted growth across adolescence and into early adulthood (Giorgio et al., 2008; Tamnes et al., 2010). Structural abnormalities, such as reduced grey matter volume in the mPFC and weaker integrity of the uncinate fasciculus, have been reported in adults with anxious temperament and anxiety disorders (Baur, Hänggi, & Jäncke, 2012; Kim & Whalen, 2009; Phan et al., 2009; Shang et al., 2014; Tromp et al., 2012). While the number of studies examining structural changes in anxious vs. non-anxious adolescents is limited, there is some evidence of similar structural abnormalities in ventral portions of the prefrontal cortex and the uncinate fasciculus for anxious youth (Liao et al., 2014; Mueller et al., 2013; Newman et al., 2015; Strawn et al., 2015; Strawn, Wehry, et al., 2013). Findings in the amygdala regions for anxious vs. non-anxious adolescents are less clear, with some studies reporting larger volumes in the right and left amygdala, specifically the basolateral amygdala (De Bellis et al., 2000; Qin et al., 2014), smaller volumes in the left amygdala (Blackmon et al., 2011; Milham et al., 2005; Mueller et al., 2013) or no difference in amygdala volume (Strawn, Chu, et al., 2013).

The functional and structural evidence outlined above, combined with the frequently reported emergence of phobic anxiety disorders during late childhood and adolescence (de Lijster et al., 2016), suggest this period to be an important and vulnerable window of fear extinction circuitry development. Consequently, given the on-going development of fear extinction circuitry, adolescents may be less responsive to traditional treatments for anxiety disorders that are based on fear extinction models, such as exposure therapy (Cartwright-Hatton, Roberts, Chitsabesan, Fothergill, & Harrington, 2004). Therefore, examining the function and structure of fear extinction circuitry in healthy adolescents may not only further our understanding of how fear extinction processes develop, but it may shed light on how treatment strategies can be tailored to the neurodevelopmental profile of vulnerability to anxiety.

In the current study we examined: (1) age-dependent changes in recruitment of the amygdala and mPFC during fear extinction, and (2) the age-dependent relationship between structure and function in the amygdala and mPFC. We used event-related functional magnetic resonance imaging (fMRI) of a cued fear conditioning paradigm, where an aversive sound served as an unconditioned stimulus and visual shapes as conditioned stimuli. In vulnerable populations, such as children and adolescents, sound stimuli are recommended over mild electric shocks in fear conditioning paradigms (Johnson & Casey, 2015; Lau et al., 2011; Pattwell et al., 2011; Pattwell et al., 2012). CS-US pairings were 100% reinforced during fear acquisition to match the reinforcement schedule typically employed in the developmental animal literature (Kim, Hamlin, & Richardson, 2009; Kim, Li, & Richardson, 2011; McCallum, Kim, & Richardson, 2010; Pattwell, Bath, Casey, Ninan, & Lee, 2011; Pattwell et al., 2012). We have previously shown the same experimental design to successfully evoke concomitant fear extinction psychophysiology and brain activation in adult populations (Morriss, Christakou, & Van Reekum, 2015, 2016), similar to other labs using 100% reinforcement (Dunsmoor, Bandettini, & Knight, 2007). In addition, we collected structural magnetic resonance imaging (sMRI) and diffusion tensor imaging (DTI) data to measure structural changes in grey matter in the mPFC and amygdala, and in white matter in the uncinate fasciculus respectively.

We hypothesized that younger adolescents would show relatively blunted fear extinction. We expected this effect to be indexed by greater amygdala activation, reduced mPFC recruitment and elevated behavioral responses (ratings) to the learned threat vs. safety cues during fear extinction. Furthermore, we hypothesised that the pattern of functional activation in adolescent participants would be associated with higher grey matter in the mPFC, lower grey matter in the amygdala, and weaker structural integrity in the uncinated fasciculus that connects the two regions.

## 1.2 Materials and Methods

### 1.2.1 Participants

55 right-handed volunteers took part in this study (M age = 17.75 yrs, SD age = 3.65 yrs, range = 12-28 yrs; 35 females & 20 males). All participants had normal or corrected to normal vision. Adult participants provided written informed consent, adolescent participants provided written informed assent and parental/guardian consent, and received a picture of their brain and £20 for their participation. The procedure was approved by the University of Reading Ethics Committee.

### 1.2.2 Procedure

Participants arrived at the laboratory and were informed of the experimental procedures. First, participants (and parents/guardians) completed consent forms as an agreement to take part in the study. Second, a hearing test was performed with an audiometer (all participants fell within the normative range of 500-8000 Hz, below 30 dB). Third, participants completed a battery of cognitive tasks (Switch Task, Stroop Task, Letter Memory Task, data not reported here) and questionnaires on a computer outside of the scanner (BISBAS, Trait Anxiety, Intolerance of Uncertainty, data not reported here). Next, participants were taken to the MRI unit. We used a conditioning task inside the scanner, whilst concurrently recording ratings (as well as pupil dilation and electrodermal activity, data not reported here). After scanning, participants rated sound stimuli presented in the scanner and completed another computerised task (not reported here).

### 1.2.3 Conditioning task

The conditioning task was designed using E-Prime 2.0 software (Psychology Software Tools Ltd, Pittsburgh, PA). Visual stimuli were presented through MRI-compatible VisualSystem head-coil mounted eye goggles (Nordic Neuro Lab, Bergen, Norway), which displayed stimuli at 60 Hz on an 800 × 600 pixel screen, with a field of view of 30° × 23°. Sound stimuli were presented through MRI-compatible AudioSystem headphones (Nordic Neuro Lab, Bergen, Norway).

Visual stimuli were light blue and yellow squares with 183 × 183 pixel dimensions that were matched for brightness. The aversive sound stimulus consisted of a fear inducing female scream from the International Affective Digitized Sound battery (IADS-2, sound number 277; (Bradley & Lang, 2007). We used Audacity 2.0.3 software (http://audacity.sourceforge.net/) to shorten the female scream to 1000 ms in length and to amplify the sound by 15 dB, resulting in a 90 dB (within +/−5 dB of 90dB) sound. An audiometer was used before testing to standardize the sound volume across participants.

The conditioning task consisted of three learning phases that were presented in three separate blocks across the scan. (1) In acquisition, one of the squares was paired with the aversive 90 dB scream (CS-), whilst the other square was presented alone (CS-). (2) In extinction, both stimuli were unpaired (CS+, CS-). (3) During reacquisition, CS+ squares were paired with the sound 25% of the time, and the CS- remained unpaired. We focus our reporting on the acquisition and extinction phases.

The acquisition phase consisted of 24 trials (12 CS+, 12 CS-), the extinction phase of 32 trials (16 CS+, 16 CS-). Experimental trials within the conditioning task were pseudorandomized into an order, which resulted in no more than three presentations of the same stimulus in a row. Conditioning contingencies were counterbalanced, with half of the participants receiving the US with a blue square and the other half of participants receiving the US with a yellow square. Participants were instructed to attend and listen to the stimulus presentations, as well as respond to a rating scale that followed each trial. The rating scale asked how ’uneasy’ the participant felt after each stimulus presentation, where the scale was 0 ’not at all’-10 ’extremely’. Participants used a hand-held MRI-compatible response box in their dominant right hand.

The presentation times of the acquisition phase were: 1500 ms square paired with a 1000 ms sound played 500 ms after the onset of a CS+ square, 3000 - 6450 ms blank screen, 4000 ms rating scale, and 1000-2500 ms blank screen. The presentation times were the same for the extinction phase, but the 1000 ms sound was omitted. To avoid predictability of stimulus on/offset, and to maximize design efficiency, we introduced jitter by randomly selecting the duration of the blank screens for each trial from the range indicated above.

### 1.2.4 Sound stimulus rating

Participants rated the valence and arousal of the sound stimulus using 9 point Likert scales ranging from 1 to 9 (Valence: negative to positive; Arousal: calm to excited).

### 1.2.5 Behavioral data scoring and reduction

Rating data were reduced for each subject by calculating their average responses for each experimental condition using the E-Data Aid tool in E-Prime (Psychology Software Tools Ltd, Pittsburgh, PA).

### 1.2.6 MRI

Participants were scanned with a 3T Siemens Trio set up with a 12 channel head coil (Siemens Inc., Erlangen, Germany). Three T2*-weighted echo planar imaging (EPI) functional scans were acquired for each phase of the conditioning task consisting of 161, 208, and 380 volumes respectively (TR = 2000 ms, TE = 30 ms, flip angle = 90°, FOV = 192 × 192 mm, 3 × 3 mm voxels, slice thickness 3 mm with an interslice gap of 1 mm, 30 axial slices, interleaved acquisition).

Following completion of the functional scans, fieldmap and structural scans were acquired, which comprised of a high-resolution T1-weighted anatomical scan (MP-RAGE, TR = 2020 ms, TE = 2.52 ms, flip angle = 90°, FOV = 256 × 256 mm, 1 × 1 × 1 mm voxels, slice thickness 1 mm, sagittal slices), two fieldmaps (TR = 488 ms, TE 1 = 4.98 ms, TE 2 = 7.38 ms, flip angle = 60°, FOV = 256 × 256 mm, slice thickness 4 mm with an interslice gap of 4 mm, 30 axial slices) and diffusion weighted images (TR = 6800ms, TE = 93 ms, flip angle = 60°, FOV = 192 × 192 mm, slice thickness 2 mm with an interslice gap of 2 mm, *b*-value =1000, 64 axial slices, 30 diffusion gradients).

### 1.2.7 fMRI analysis

FMRI analyses were carried out in Feat version 5.08 as part of FSL (FMRIB’s Software Library, www.fmrib.ox.ac.uk/fsl). Brains were extracted from their respective T1-weighted images by using the FSL Brain Extraction Tool (BET) (Smith, 2002). Distortion, slice timing and motion correction were applied to all EPI volumes using FUGUE and MCFLIRT tools (Jenkinson, Bannister, Brady, & Smith, 2002). Gaussian smoothing (FWHM 5mm) and a 50 second high-pass temporal filter were applied.

A first-level GLM analysis was carried out for each functional scan run from each learning phase. Separate regressors were specified for the experimental conditions of primary interest in each learning phase (acquisition: CS+/CS−, extinction: CS+ /CS) by convolving a binary boxcar function with an ideal haemodynamic response (HR), which corresponded to the length of each trial (1500 ms). Regressors for the uneasiness rating period, six motion parameters and any head movements above 1mm were included to model out brain activity or movement artefacts that were unrelated to the conditions of interest.

We defined two main effect contrasts to reveal fear extinction-related activity. To examine temporal effects across extinction, we contrasted (CS+ vs. CS-)_EARLY_> (CS+ vs. CS-) _LATE_. We defined early extinction as the first eight trials for CS+ and CS-and the last eight trials for CS+ and CS-. We also examined the overall effect of CS+ vs. CS- during extinction. All contrasts were normalized and registered to MNI standard space using FLIRT (Jenkinson, Bannister, Brady, & Smith, 2002). Second-level GLM analysis consisted of regressors for the group mean and a linear regressor for demeaned age scores using FSL’s Ordinary Least Squares procedure.

We were specifically interested in the extent to which age would be associated with the BOLD response in the amygdala and mPFC for fear extinction. Therefore, we performed small volume corrections on the left amygdala, right amygdala and prefrontal cortex using cluster thresholding with a z = 2.3 and a corrected p < 0.05 on the age × extinction (CS+ vs. CS-) and age × extinction (CS+ vs. CS-) _EARLY>_ (CS+ vs. CS-) _LATE_ contrast maps. We used anatomically defined masks from the Harvard-Oxford cortical and subcortical structural atlases in FSL (Desikan et al., 2006). We selected the left amygdala, right amygdala and prefrontal (including the subgenual anterior cingulate, medial and orbitofrontal portions, and excluding the lateral portion) cortical regions with a 50% probability threshold. To allow for comparisons across studies, we also carried out a whole-brain analysis using cluster thresholding with a z = 2.3 and a corrected p < 0.05, and included CS- vs. CS+.

### 1.2.8 Grey matter probability in the amygdala and mPFC

Processing of structural images was performed in FSL. Firstly, structural images were brain-extracted using BET (Smith, 2002). Secondly, structural images were segmented based on tissue-type using FMRIB’s Automated Segmentation Tool (FAST) (Zhang, Brady, & Smith, 2001). Thirdly, any resulting areas of age-related activation in the amygdala or mPFC identified from the fMRI analysis were transformed into structural space for each subject. Lastly, we extracted grey matter probability estimates of any resulting areas of age-related activation in the amygdala or mPFC identified from the fMRI analysis from each subjects’ segmented structural image.

### 1.2.9 White matter integrity of the uncinate fasciculus

Diffusion-weighted image processing in FSL included corrections for motion, eddy currents and inhomogeneities in the magnetic field. Then the tensor model was fitted using FDT (FMRIBS Diffusion Toolbox) in order to calculate FA values for each voxel, producing one FA image per subject. Voxels with FA values lower than 0.2 were removed. We created 25% probability masks of the left and right uncinate fasciculus the forceps minor and the corticospinal tract from the JHU white-matter tractography atlas (Mori, Wakana, Van Zijl, & Nagae-Poetscher, 2005). The forceps minor and corticospinal tract served as control regions, similar to previous structure-function research (Swartz, Carrasco, Wiggins, Thomason, & Monk, 2014). All tract masks were transformed into diffusion space and applied to each subjects’ FA image, resulting in an FA value for each tract per subject.

### 1.2.10 Statistical analyses

Main effects of conditioning and age in fear extinction were assessed by conducting a Condition (CS+, CS-) × Time (Early, Late) × Age (days) repeated measures ANCOVA on behavioral ratings. The early part of extinction was defined as the first eight CS+ and CS- trials, and the last part of extinction was defined as the last eight CS+ and CS- trials.

We correlated any resulting areas of age-related activation in the amygdala or mPFC identified from the fMRI analysis with grey matter probability estimates of those regions and FA values in the uncinate fasciculus. To examine whether the relationship between mPFC activation during extinction and FA values was specific to the uncinate fasciculus we conducted control analyses in the forceps minor and corticospinal tract.

Furthermore, if there were relationships between age and function/structure then we conducted follow up mediation analyses to assess whether structure mediated the relationship between age and functional engagement of the amygdala or mPFC. We used Freedman-Schatzkin mediation tests, where the difference in the regression coefficient between the unadjusted association between the independent variable and the dependent variable is compared with the regression coefficient for this association when it is adjusted for a potential intervening or mediating variable (Freedman & Schatzkin, 1992).

## 1.3 Results

Three participants did not complete the scanning procedure and three participants were removed due to excessive head movements, leaving forty-nine participants for analysis (M age = 18.70 yrs, SD age = 3.64 yrs, range = 12-28 yrs; 31 females & 18 males).

### 1.3.1 Ratings

All participants rated the sound stimulus as aversive (M = 2.96, SD = 1.17) and moderately arousing (M = 5.10, SD = 1.94). Sound arousal ratings were negatively correlated with age, *r*(47) = −.286, *p* = .047 such that increasing age was associated with lower rated arousal. Sound valence ratings did not correlate with age, *r*(47) =-.141, *p* = .333.

During extinction, participants reported feeling significantly more uneasy during the CS+ (M = 1.04, SD = 1.16) compared to the CS- (M = 0.92, SD = 1.14) trials across the extinction phase, *F*(1,47) = 5.094, *p* = .029, η^2^ = .098. In addition, participants also reported feeling more uneasy at the start of extinction, compared to the end of extinction *F*(1,47) = 6.875, *p* = .012, η^2^ = .128. Contrary to predictions, there was no interaction of Condition × Time, *F*(1,47) = 1.004, *p* = .322 for the uneasiness ratings.

Results revealed no age differences for uneasiness ratings in any of the experimental phases, max *F* =.815.

### 1.3.2 fMRI

A full list of details of clusters of activation in a-priori regions of interest and whole-brain corrected for key contrasts is provided in Table 1. During fear extinction we found a significant interaction between Condition × Age in right amygdala, such that younger age predicted greater right amygdala activity to the CS+ vs. CS- (see Fig 1 and Table 1)^1^.

**Figure 1.**
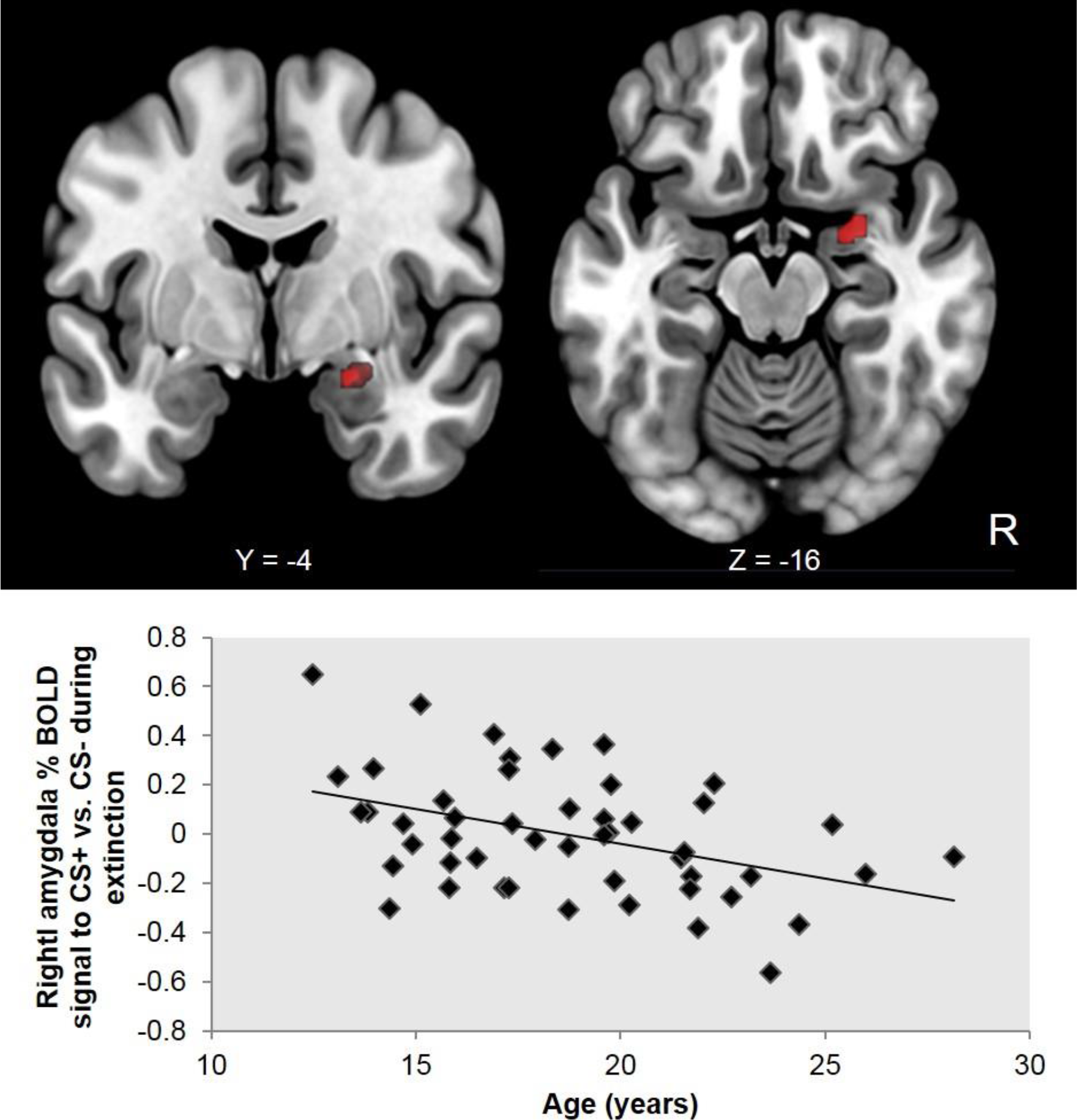
Top panel: Right amygdala activation to CS+ > CS- × decreasing age. Bottom panel: Younger age is significantly associated with greater right amygdala activation to the CS+ vs. CS- across the fear extinction phase. Age, measured in days (age is in years for display purposes only). Coordinates in the top panel are in MNI space. R=right.

**Table 1.** Regional activation patterns in response to extinction

| Contrast | Brain region | BA | Voxels | Max Z | Location of max Z |  |  |
| --- | --- | --- | --- | --- | --- | --- | --- |
|  |  |  |  |  | x | y | z |
| **Age effects** |  |  |  |  |  |  |  |
| CS+ > CS- x decreasing age | A-priori ROI/SVC: right amygdala |  | 20 | 3.1 | 26 | 0 | -16 |
| CS+ > CS- x decreasing age | R putamen, R amygdala, R hippocampus |  | 281 | 3.28 | 20 | -90 | 32 |
| CS+ > CS- x decreasing age | R lateral occipital cortex, R cuneal cortex |  | 271 | 3.6 | 14 | -2 | -22 |
| CS+ > CS- x increasing age | L postcentral gyrus |  | 423 | 3.87 | -54 | -22 | 50 |
| CS+ > CS- early - CS+ > CS- late x increasing age | A-priori ROI/SVC: medial frontal cortex, paracingulate gyrus, frontal pole |  | 174 | 3.89 | 0 | 66 | 10 |
| CS+ > CS- early - CS+ > CS- late x increasing age | L frontal pole |  | 313 | 3.92 | -22 | 42 | 50 |
| **Main effects** |  |  |  |  |  |  |  |
| CS - CS+ | Posterior cingulate, precuneus cortex, L intracalcarine cortex, L lingual gyrus |  | 1365 | 4 | -24 | -42 | -14 |
| CS - CS+ | L postcentral gyrus, L precentral gyrus |  | 502 | 4.74 | -48 | -14 | 48 |
| CS - CS+ | L postcentral gyrus, L precentral gyrus |  | 392 | 3.38 | -24 | -52 | 74 |
| CS - CS+ | R lateral occipital cortex |  | 387 | 4.35 | 42 | -64 | 26 |
| CS - CS+ | L lateral occipital cortex |  | 290 | 3.55 | -42 | -76 | 40 |
| CS - CS+ | L lingual gyrus |  | 274 | 3.85 | -22 | -60 | -14 |
| CS - CS+ | R postcentral gyrus, R precentral gyrus |  | 256 | 4.22 | 40 | -34 | 32 |
| CS+ > CS- early - CS+ > CS- late | A-priori ROI/SVC: left orbital frontal cortex |  | 200 | 3.81 | -34 | 34 | -14 |
| CS+ > CS- early - CS+ > CS- late | L occitpital pole |  | 383 | 5.02 | 28 | -96 | -4 |
| CS+ > CS- early - CS+ > CS- late | R occipital pole |  | 277 | 3.86 | -34 | -98 | -4 |
| CS- > CS+ early - CS- > CS+ late | R postcentral gyrus, R supramarginal gyrus |  | 419 | 4.01 | 52 | -20 | 36 |
| CS- > CS+ early - CS- > CS+ late | L postcentral gyrus, L supramarginal gyrus |  | 378 | 3.75 | -50 | -28 | 38 |
| CS- > CS+ early - CS- > CS+ late | L lingual gyrus |  | 359 | 4.27 | -16 | -68 | -8 |

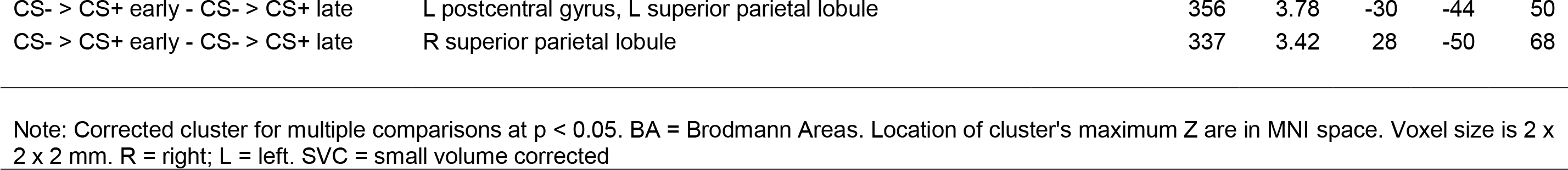

From the analysis of extinction over time, we found a significant interaction between Condition × Time × Age in the mPFC. Older age was associated with stronger recruitment of the mPFC to CS+ vs. CS- in the early trials, and younger age was associated with stronger recruitment of the mPFC to the CS+ vs. CS- late trials (see Fig 2 and Table 1).

**Figure 2.**
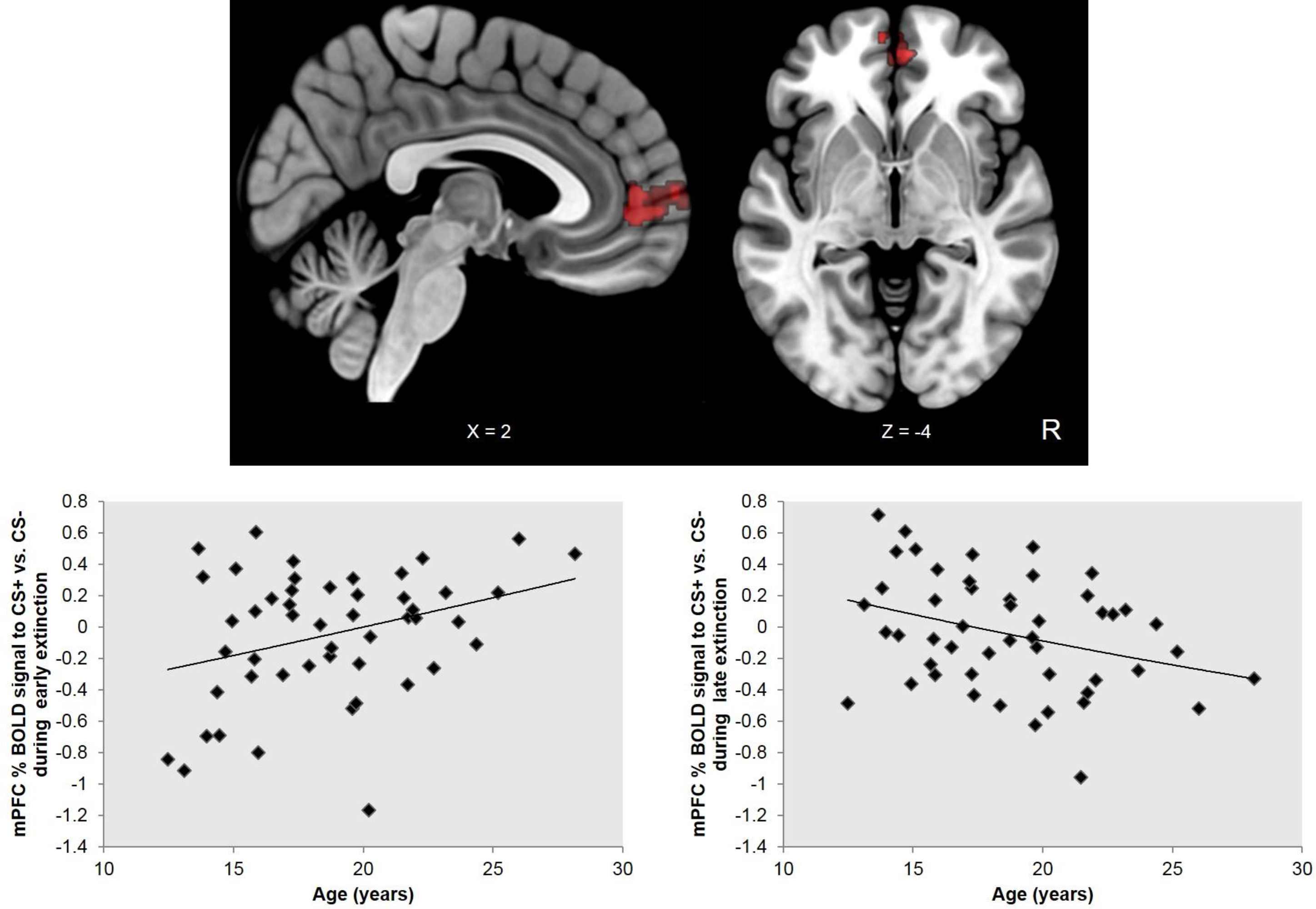
Top panel: mPFC activation to CS+ > CS- early - CS+ > CS- late × increasing age. Bottom left and right panels: Older age is significantly associated with greater mPFC activity to the CS+ vs. CS- during early fear extinction, whilst younger age is significantly associated with greater mPFC activity to the CS+ vs. CS- during late fear extinction. Age, measured in days (age is in years for display purposes only). Coordinates in the top panel are in MNI space. R=right.

### 1.3.3 Grey matter probability in the amygdala and mPFC

As mentioned in the Methods section, we extracted grey matter probability from the fMRI clusters identified in a-priori regions of interest and depicted in Fig 1 & 2, i.e. right amygdala and mPFC.

#### Age effects

As expected, we found a significant negative correlation between mPFC grey matter probability and age, *r*(47) = −.436, *p* = .002. No significant correlation between the right amygdala and age was observed, *r*(47) = .139, *p* = .340.

#### Structure-Function relationships

Grey matter probability in the mPFC significantly predicted the mPFC activation difference score for extinction (CS+ vs. CS _−Early_ − CS+ vs. CS _−Late_), *r*(47) = −.342, *p* = .016. We found no significant relationships between grey matter probability in the mPFC or right amygdala with right amygdala activation during extinction, *r*(47) = .018, *p* = .904; *r*(47) = −.167, *p* = .250.

#### Mediation by structure

Grey matter probability in the mPFC did not significantly mediate the relationship between age and mPFC activity during extinction, *t* = 1.103, *p* = .275.

### 1.3.4 White matter integrity of the uncinate fasciculus

#### Age effects

In line with predictions, we found positive correlations with age and structural integrity of the bilateral uncinate fasciculus (the left and right uncinate fasciculus FA values significantly correlated, *r*(47) = .73, *p* < .001, thus we collapsed across hemisphere), *r*(47) = .30, *p* = .035, suggesting increased white matter integrity of this tract with advancing age.

#### Structure-Function relationships

Greater structural integrity of the bilateral uncinate fasciculus predicted the mPFC activity difference score for extinction (CS+ vs. CS _−Early_ − CS+ vs. CS _−Late_), *r*(47) = .463, *p* = .001. As highlighted in the Introduction, white matter develops across the brain during adolescence. The bilateral uncinate fasiculus significantly predicted mPFC activity during extinction (second step: Δ*R*^2^=.173, *F*(1,45) = 10.367, *p* = .002), over and above the forceps minor and corticospinal tract (first step: *R*^2^=.078, *F*(2,46) = 1.933, *p* = .156).

Structural integrity of the bilateral uncinate fasciculus was not significantly associated with right amygdala activity for CS+ vs. CS- during extinction, *r*(47) = −.066, *p* = .655.

#### Mediation by structure

The structural integrity of the bilateral uncinate fasciculus significantly mediated the relationship between age and mPFC activity during extinction (difference score: CS+ vs. CS _−Early_ − CS+ vs. CS _−Late_), *t* = 2.695, *p* = .009 (see Fig 3). In the mediation model, after adjustment for the bilateral uncinate fasciculus, age continued to predict mPFC activity but to a lesser extent (see Fig 3).

**Figure 3.**
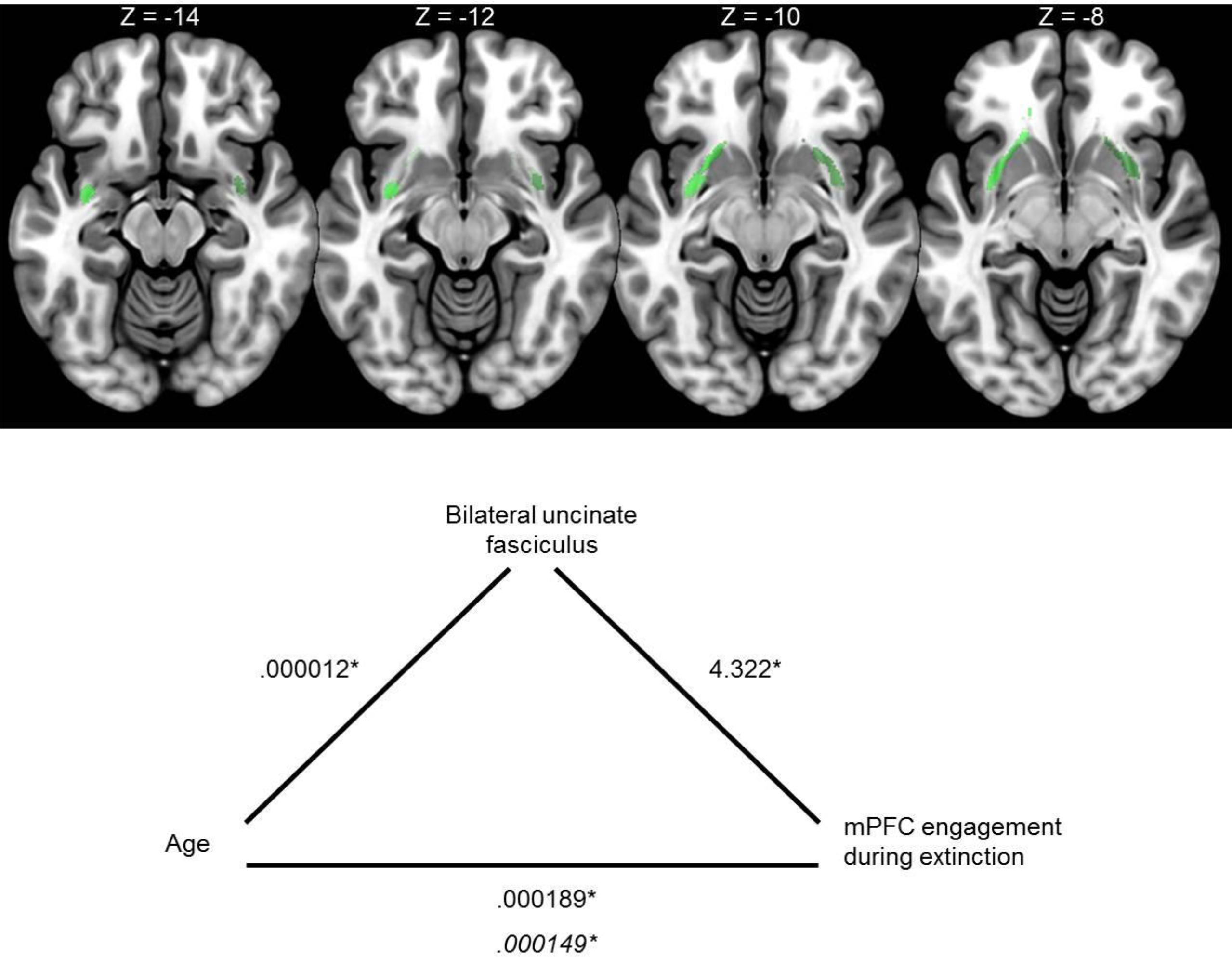
Top panel: Bilateral uncinate fasciculus brain masks. Bottom panel: Greater structural integrity of the unincate fasciculus mediated the relationship between age and mPFC engagement during extinction (difference score: CS+ vs. CS _−Early_ − CS+ vs. CS _−Late_). Coordinates in the top panel are in MNI space. Values in the bottom panel represent the unstandardized coefficients from linear regression models. Italicized values represent the unstandardized coefficient for age and mPFC engagement after adjustment for the uncintate fasciculus mediator. *p < .05

## 1.4 Discussion

In the current study, we show that amygdala and mPFC recruitment during fear extinction matures during adolescence and into young adulthood. Our data suggest that adolescence is associated with blunted fear extinction, through delayed recruitment of the mPFC and prolonged engagement of the amygdala to threat cues compared to safety cues. Furthermore, the relationship between age and delayed recruitment of the mPFC was mediated in part by the structural integrity of the uncinate fasciculus, a white matter tract connecting the amygdala and mPFC. These data contribute to the growing developmental literature, describing adolescence as a vulnerable period of development, with increased risk to anxiety disorders (Casey, Oliveri, & Insel, 2014; Giedd, Keshavan, & Paus, 2008).

During fear extinction, younger age was characterized by increased amygdala activity to threat vs. safety cues, consistent with previous rodent and human fear extinction studies (Johnson & Casey, 2015; Kim, Li, & Richardson, 2011; Pattwell, Bath, Casey, Ninan, & Lee, 2011; Pattwell et al., 2012), suggesting exaggerated fear expression during adolescence. Furthermore, younger age was associated with delayed mPFC activity to threat vs. safety cues during extinction, in line with previous rodent work (Kim, Li, & Richardson, 2011; Pattwell, Bath, Casey, Ninan, & Lee, 2011; Pattwell et al., 2012), suggesting compromised fear regulation during adolescence. These data suggest that adolescents may show less efficient fear extinction, rendering them less responsive to current exposure-based therapies for anxiety. A recent study by Johnson et al. (2015) has shown that compromised fear extinction in adolescents may be aided with memory reconsolidation techniques, which remind adolescents of safety, subsequently blocking the recovery of learned fear.

Our data also show structural maturation in fear extinction circuitry. In line with prior structural work (Giedd, 2004; Gogtay et al., 2004; Lebel & Beaulieu, 2011; Lebel, Walker, Leemans, Phillips, & Beaulieu, 2008; Østby et al., 2009; Tamnes et al., 2010; Wierenga et al., 2014), grey matter in the mPFC decreased with age, while white matter in the uncinated fasciculus increased with age. Structural maturational changes in the mPFC and uncinate fasciculus were associated with age-dependent changes in mPFC responding during fear extinction. Grey matter in the amygdala was not correlated with amygdala BOLD response during extinction.

However, only the uncinate fasiculus was found to mediate the relationship between age and mPFC responding during fear extinction. After adjustment for the contribution of the uncinate fasciculus in the mediation model, age continued to predict mPFC activity but to a lesser extent. These results suggest that both biological age and structural changes in the uncinate fasciculus contribute to mPFC engagement during extinction. Furthermore, we can speculate that white matter integrity of the uncinate fasciculus may be a better marker of developmentally driven blunted fear extinction, over and above grey matter pruning of the mPFC (and grey matter growth in the amygdala).

Notably, weaker structural integrity of the uncinate fasciculus has been found in populations with anxious temperament (Baur, Hänggi, & Jäncke, 2012; Kim & Whalen, 2009; Liao et al., 2014; Phan et al., 2009; Tromp et al., 2012), suggesting that this tract may play a crucial role in anxiety aetiology. The uncinate faciculus has a protracted growth, as it continues to develop until the third decade of life, unlike the expedited grey matter pruning in regions such as the mPFC. On this basis, following the development of the uncinate faciculus may be particularly promising for neurodevelopmental researchers and clinicians. Future work with longitudinal designs may be able to further elucidate the relationship between fear extinction ability, white matter integrity in the uncinate fasciculus and anxiety disorder vulnerability by tracking the extent of malleability within the uncinate fasciculus across adolescence and into early adulthood. From this we may be able to identify whether anxious individuals have a different developmental trajectory of the uncinate faciculus and when it may be best to attempt treatment or intervention.

To conclude, we describe age-dependent relative blunting of fear extinction during adolescence. In our sample, younger age was associated with exaggerated amygdala responses to learned threat vs. safety cues during fear extinction. In addition, younger age was associated with reduced mPFC activity during early extinction. The relationship between age and delayed recruitment of the mPFC was mediated by the structural integrity of the uncinate fasciculus, a white matter tract connecting the amygdala and mPFC. These findings suggest reduced flexibility in amygdala and mPFC during adolescence. Importantly, these results highlight the need for further examination of structural and functional changes in amygdala-mPFC circuitry during adolescence, with respect to: (1) risk of anxiety disorder development, (2) effectiveness of current exposure based therapies, and (3) development of novel anxiety disorder treatments that are age specific and age appropriate (Casey, Glatt, & Lee, 2015; Casey, Oliveri, & Insel, 2014; Johnson & Casey, 2015).

## Footnotes

1 In addition, we observed a homologous area in left amygdala where younger age predicted greater left amygdala activity to the CS+ vs. CS-, but this cluster did not survive corrections for multiple comparisons.

